# Neural tracking of the fundamental frequency of the voice: the effect of voice characteristics

**DOI:** 10.1101/2020.08.26.267922

**Authors:** Jana Van Canneyt, Jan Wouters, Tom Francart

**Affiliations:** ExpORL, Dept. of Neurosciences, KU Leuven, Herestraat 49 bus 721, 3000 Leuven, Belgium

**Author notes:** Email addresses:* (Jana Van Canneyt), (Jan Wouters), (Tom Francart).

**Keywords:** f0 tracking, natural speech decoding, auditory processing, stimulus parameters, pitch processing

## Abstract

Traditional electrophysiological methods to study temporal auditory processing of the fundamental frequency of the voice (f0) often use unnaturally repetitive stimuli. In this study, we investigated f0 processing of meaningful continuous speech. EEG responses evoked by stories in quiet were analysed with a novel method based on linear modelling that characterizes the neural tracking of the f0. We studied both the strength and the spatio-temporal properties of the f0-tracking response. Moreover, different samples of continuous speech (six stories by four speakers: two male and two female) were used to investigate the effect of voice characteristics on the f0 response.

The results indicated that response strength is inversely related to f0 frequency and rate of f0 change throughout the story. As a result, the male-narrated stories in this study (low and steady f0) evoked stronger f0-tracking compared to female-narrated stories (high and variable f0), for which many responses were not significant. The spatio-temporal analysis revealed that f0-tracking response generators were not fixed in the brainstem but were voice-dependent as well. Voices with high and variable f0 evoked subcortically-dominated responses with a latency between 7 and 12 ms. Voices with low and steady f0 evoked responses that are both subcortically (latency of 13-15 ms) and cortically (latency of 23-26 ms) generated, with the right primary auditory cortex as a likely cortical source. Finally, additional experiments revealed that response strength greatly improves for voices with strong higher harmonics, which is particularly useful to boost the small responses evoked by voices with high f0.

## 1. Introduction

Temporal processing of the fundamental frequency of the voice (f0) is often studied with event-related auditory-evoked potentials, usually envelope following responses (EFRs) (e.g. Krishnan et al. (2004); Peng et al. (2018); Van Canneyt et al. (2020)). Typically, a short stimulus is presented hundreds of times while measuring the phase-locked neural responses to the f0 with EEG or MEG. The large amount of repetitions is needed to increase the signal to noise ratio (SNR) by averaging out the response trials. These classic EFR paradigms have proven their worth but the high degree of repetition and predictability in the evoking stimulus fundamentally impacts auditory processing and perception. Continuous speech stimuli better approximate natural language use and as a result, research findings with these stimuli are more relevant for auditory processing in day-to-day communication (Theunissen et al., 2000). Moreover, the continuous speech stimuli are often stories, which increases the motivation and attention of the subjects (Hamilton and Huth, 2020). This is particularly interesting for clinical populations that are harder to test, e.g. children or people with a mental disability. A final benefit of using continuous speech is that the evoked responses reflect all aspects of speech processing, from fundamental acoustic processing to higher-order lexical and semantic processing, and all these speech processing stages can be studied with the same data (Brodbeck and Simon, 2020). Inspired by these advantages, we investigated the temporal processing of the f0 using continuous speech stimuli.

To process neural responses to the f0 in continuous speech, we used a framework based on linear decoding and encoding models developed throughout the last decade (e.g. Mesgarani et al. (2009); Lalor and Foxe (2010); Ding and Simon (2012); Crosse et al. (2016); Vanthornhout et al. (2018)). A linear decoding model, or backward model, reconstructs a given stimulus-related feature from a linear combination of multi-channel neural responses and their time-lagged versions (Mesgarani et al., 2009). The model is usually trained on part of the data (e.g. 80 % of the available data) and then tested on the remainder of the data. The correlation between the reconstructed feature and the actual feature is used to evaluate the quality of the model and is related to response strength. The advantage of a backward model is that it employs information from multiple recording channels simultaneously, but the disadvantage is that the weights of the backward model cannot be interpreted (Haufe et al., 2014). Conversely, a linear encoding model, or a forward model, reconstructs the neural response in each recording channel separately, based on a given stimulus-related feature and time-lagged versions of it. The set of weights needed to predict one channel of the neural response from the feature is called a temporal response function (TRF) (Ding and Simon, 2012) and, in contrast with the backward model, these sets of weights (or TRFs) can be interpreted. They can be understood as the impulse response of the auditory system and provide both spatial and temporal information about the response.

These linear models can be constructed for various stimulus features and depending on the feature, different aspects of auditory processing can be targetted. The most commonly used feature is the speech envelope (envelope-tracking), which targets relatively low-level cortical speech processing (e.g. Ding and Simon (2014); Vanthornhout et al. (2018)). Other features target higher-order cortical processing, e.g. phoneme and phonetic representations, the spectrogram, entropy, phoneme surprisal or semantic dissimilarity (Di Liberto et al., 2015; Di Liberto and Lalor, 2017; Di Liberto et al., 2018; Brodbeck et al., 2018; Broderick et al., 2018; Lesenfants et al., 2019a,b). Responses to the above mentioned features, and particularly responses to the speech envelope, have been studied quite extensively in recent years (e.g. Etard and Reichenbach (2019); Verschueren et al. (2019); Decruy et al. (2019, 2020)). In contrast, research that analyses continuous responses with an f0-based feature (f0-tracking), targetting subcortical acoustic processing, is just emerging: Forte et al. (2017) developed a method to process continuous f0 responses based on crosscorrelation. In continuation, Etard et al. (2019) performed attention decoding experiments with f0-tracking through linear modelling. Finally, Saiz-Alia and Reichenbach (2020) used the same dataset to simulate continuous f0 responses with a model of the auditory system. Due to the natural and engaging stimulus, f0-tracking has large clinical potential. However, this novel method needs to be characterized in normal hearing young adults, before its applications can be further explored. The current study aimed to take the next steps in this characterisation.

At this time, only a very limited amount of information is available on f0-tracking. In contrast, the classic EFR (typpically evoked by repeating short stimuli) has recently been characterised quite extensively (e.g. Purcell et al. (2004); Bidelman (2018); Van Canneyt et al. (2019, 2020)). Since the EFR and f0-tracking are both based on phaselocking to the f0, similarity between response properties is expected. However, because the continuous stimuli used for f0)tracking are more meaningful, more variable and less predictable than the typical EFR stimulus, some differences may occur. With this in mind, the present study aimed to establish a frame of reference on response strength and spatiotemporal characteristics of the continuous f0 response, while evaluating whether two important recent findings from EFR studies generalize to continuous f0 responses. The first finding concerns the neural generator(s) of the responses. In recent years, new evidence indicating cortical contributions to the EFR (Coffey et al., 2016, 2019; Bidelman, 2018) has challenged the long-standing belief that the EFR is a brainstem response, generated by the colliculus inferior (e.g. Sohmer et al. (1977); Aiken and Picton (2008)). Through TRF analysis, we aim to investigate the neural sources of f0-tracking and compare it to what has been found for classic EFRs. Cortical contributions to f0-tracking are not unlikely as the more meaningful and less predictable stimuli could solicit more top-down processing than the traditional EFR stimuli. The second finding is that higher fundamental frequencies and higher rates of fundamental frequency change in the stimulus result in smaller EFRs (Purcell et al., 2004; Billings et al., 2019; Van Canneyt et al., 2020). We aim to investigate whether similar effects are likely to occur for f0-tracking. In summary, the goals of this study are to characterise the strength and spatio-temporal properties of the continuous f0 response (including an estimation of its sources) and to investigate how f0-tracking is influenced by important voice characteristics like f0 frequency and rate of f0 change.

## 2. Methods

### 2.1. Dataset and subjects

The EEG data analysed in this paper is part of a large existing dataset (Accou et al., 2020; Monesi et al., 2020) that contains EEG responses to continuous Flemish speech for young normal hearing participants (not publicly available). A subset of 46 (41 females, 5 males) subjects was selected based on presentation with the same experimental conditions. The age of the subjects ranged between 18 and 25 years old (mean = 20.7 years, standard deviation = 1.7 years). All participants had normal hearing (all thresholds < 20 dB HL), verified using pure-tone audiometry (octave frequencies between 125 and 8000 Hz) and were native Flemish (or Dutch) speakers. The experiments were approved by the medical ethics committee of the University Hospital of Leuven and all subjects signed an informed consent form before participating (s57102).

### 2.2. Stimuli

The stimuli presented to the participants were six fairy tales with titles: *Milan, Eline, Anna en de vorst* (AEDV), *De wilde zwanen* (DWZ), *De oude lantaarn* (DOL) and *De kleine zeemeermin* (DKZ). Table 1 summarises important information about the stories. Note that *Eline, Milan* and *DOL* were presented and analysed in full, whereas the stories *AEDV, DWZ* and *DKZ* were presented in multiple sections, of which only the first section was analysed for this study. Moreover, in the original dataset two story sections, randomly selected per subject, were presented in noise and these data points were not included (leading to an uneven amount of subjects per story). However, since these data points are missing nearly completely at random, the response analysis remains largely unbiased.

**Table 1:**
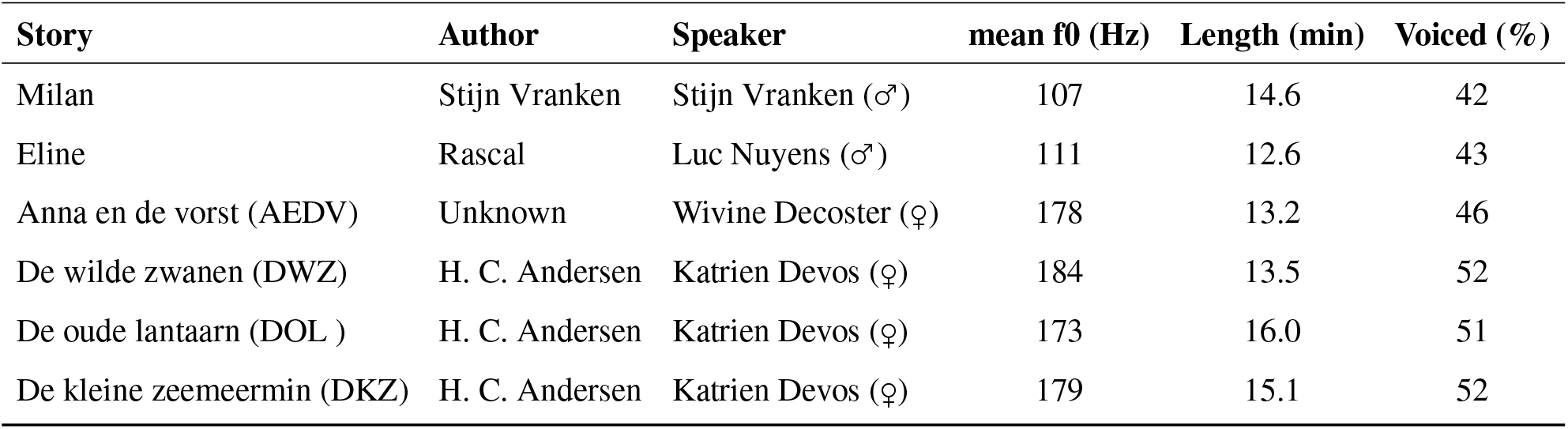
Information on the stories presented to the subjects.

| Story | Author | Speaker | mean f0 (Hz) | Length (min) | Voiced (%) |
| --- | --- | --- | --- | --- | --- |
| Milan | Stijn Vranken | Stijn Vranken (♂) | 107 | 14.6 | 42 |
| Eline | Rascal | Luc Nuyens (♂) | 111 | 12.6 | 43 |
| Anna en de vorst (AEDV) | Unknown | Wivine Decoster (♀) | 178 | 13.2 | 46 |
| De wilde zwanen (DWZ) | H. C. Andersen | Katrien Devos (♀) | 184 | 13.5 | 52 |
| De oude lantaarn (DOL) | H. C. Andersen | Katrien Devos (♀) | 173 | 16.0 | 51 |
| De kleine zeemeermin (DKZ) | H. C. Andersen | Katrien Devos (♀) | 179 | 15.1 | 52 |

### 2.3. The f0 feature

The response analysis required an f0 feature, i.e. a waveform oscillating at the instantaneous f0 of the stimulus. This f0 feature was obtained by bandpass filtering the stimulus (Etard et al., 2019) with a Chebyshev filter with 80 dB attenuation at 10 % outside the passband and a pass band ripple of 1 dB. The filter cut-offs were based on the distribution of the f0 in the story (visualised in Figure 2): the stimulus was filtered between 75 and 175 Hz for male-narrated stories and between 120 and 300 Hz for female-narrated stories. Unvoiced and silent sections of a story have no f0 and were therefore excluded from the data analysis. The technique proposed by Forte et al. (2017) allowed to determine these sections in an automatic way: parts where the envelope amplitude was less than 10% of the maximum found in the speech, were considered silent. Parts where the instantaneous f0 (determined through autocorrelation) was lower than 60 Hz, higher than 400 Hz or changed more than 10 Hz within 1 ms were considered unvoiced. The silent and unvoiced sections were removed from the f0 feature, after which the feature was normalized to have zero mean and a variance of 1. It is important to note that only about 50 % of each story was non-silent and voiced (see Table 1 for the exact percentage per story). As a consequence, only about half of the measured data could be used for the analysis.

**Figure 1:**
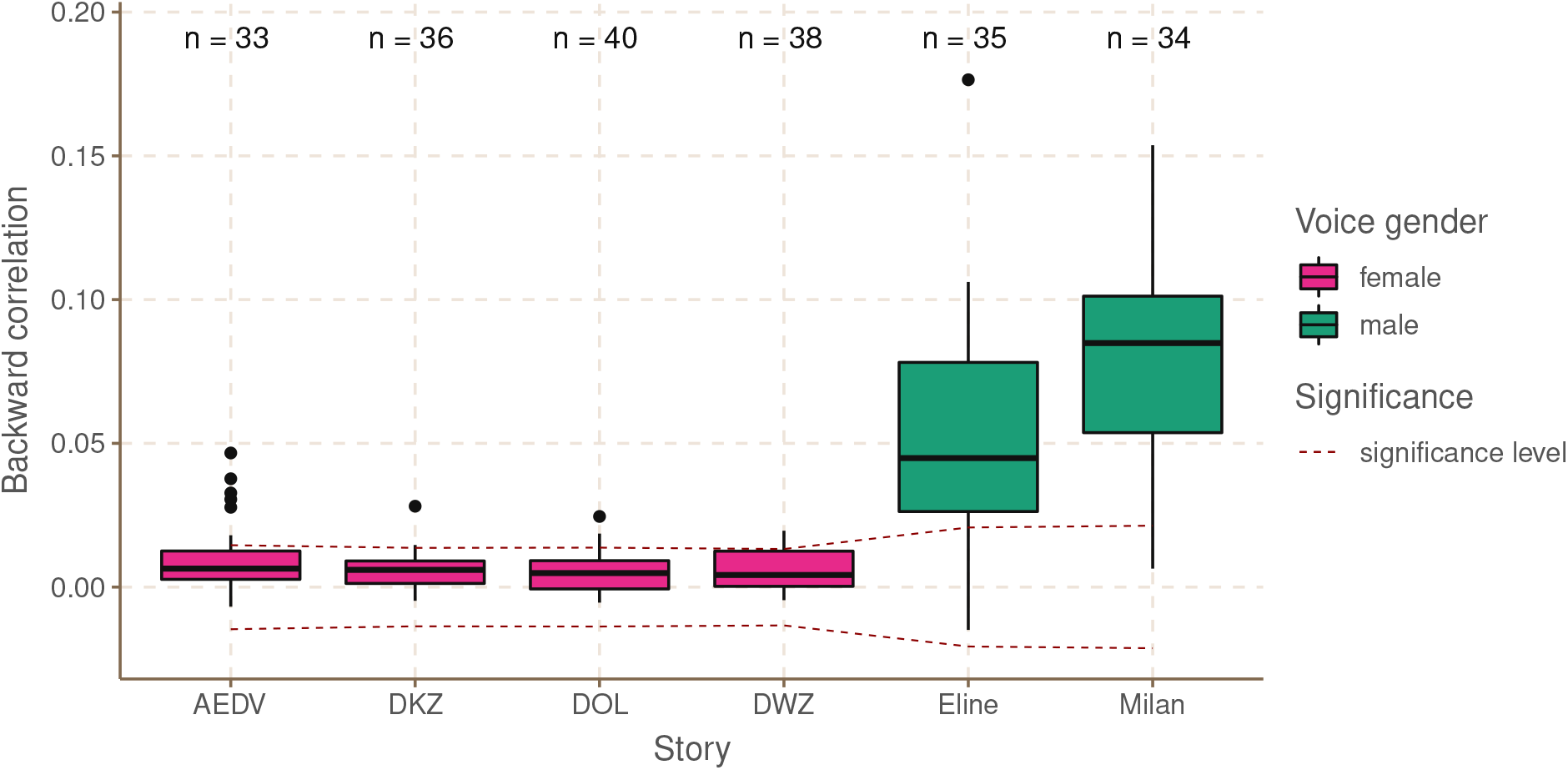
Backward correlations per story. The significance level is indicated with a dashed line. “n” denotes the number of data points in each boxplot. The colors indicate the voice gender of the speaker.

**Figure 2:**
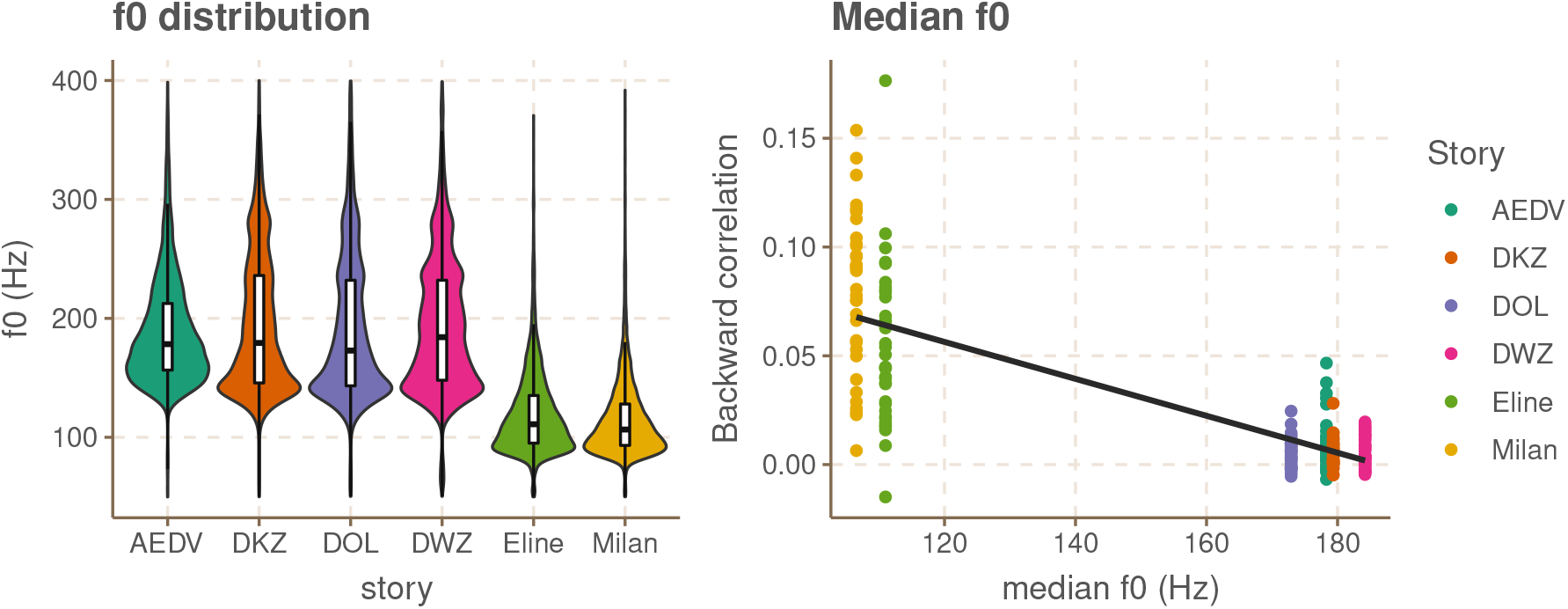
f0 distribution of the stories and the relation between backward correlations and the median f0 of the story.

### 2.4. f0-related stimulus parameters

One of the aims of this study was to investigate how f0-tracking is influenced by the f0 of the voice and the rate at which the f0 changes throughout the story. To determine the f0 distribution of each story, we estimated the f0 in 16 ms intervals across all voiced sections of the story using PRAAT (Boersma and Weenink, 2015). The derivative of the f0 estimations was taken to obtain the rate of f0 change expressed in Hz/s. The effect of median f0 and median rate of f0 change on the continuous f0 response was statistically evaluated in R (version 3.6.3., R Core Team (2018)), using linear mixed models (package lme4, version 1.1.21, Douglas et al. (2015)) with a random intercept per subject.

#### 2.4.1. EEG responses

The EEG responses in the dataset were recorded with a 64-channel Biosemi ActiveTwo EEG recording system (sampled at 8192 Hz). The 64 Ag/AgCl active scalp electrodes were placed on a cap according to the international standardized 10-10 system (American Clinical Neurophysiology Society, 2006). Two extra electrodes on the cap, CMS and DRL, functioned as the common electrode and the current return path, respectively. Subjects were seated in an electromagnetically-shielded sound-proof booth and instructed to listen carefully to the stories. These were presented binaurally and in random order through electrically-shielded insert phones (Etymotic ER-3A, Etymotic Research, Inc., IL, USA) using the APEX 3 software platform developed at ExpORL (Dept. Neurosciences, KU Leuven) (Francart et al., 2008). Stimulus intensity was set to 62 dB A in each ear. The setup was calibrated in a 2-cm^3^ coupler (Bruüel & Kjaer, type 4152, Nærum, Denmark) using stationary speech weighted noise with the same spectrum as each of the different stories. To encourage attentive listening, participants answered a question about the content of each story after its presentation.

We applied several preprocessing steps to the raw EEG-data from the dataset. First, the data was downsampled to a sampling frequency of 1024 Hz. Then, artefacts were removed using a multi-channel Wiener filter algorithm with delays from −3 to 3 samples included and a noise weighting factor of 1 (Somers et al., 2018). The data was re-referenced to the average of all electrodes and filtered with the same filter used to obtain the f0 feature (see above). We also applied a notch filter to remove the artefact caused by the third harmonic of the utility frequency at 150 Hz (the other infected frequencies did not fall in the bandpass filter range). Finally, the unvoiced and silent sections, as determined based on the stimulus, were removed and the EEG was normalized to be zero mean with unit variance.

### 2.5. Backward modelling

The EEG responses were analysed with both backward decoding and forward encoding models. Both of these were implemented in MATLAB R2016b (The MathWorks Inc., 2016) using custom scripts and the mTRF toolbox (Crosse et al., 2016). The methods for the backward and forward modelling are discussed in this section and the next section, respectively. A more detailed description of some aspects is available in the appendix.

In backward modelling, one attempts to reconstruct a known stimulus-related feature, in this case the f0 feature, based on a linear combination of the time-shifted data from the EEG electrodes. In this study, time shifts between 0-25 ms and 0-40 ms were considered for female and male voices respectively, with 1/fs steps (fs = 1024 Hz). These lags were determined based on the results of forward modelling, see Figure 4. Regularization was done using ridge regression (Tikhonov and Arsenin, 1977; Hastie et al., 2001; Machens et al., 2004). The model was estimated based on a training dataset and then tested by using that same model to predict the feature from an unseen set of EEG data. The testing was done on a section of two minutes and the training was done on the remainder of the data. To evaluate the performance of the model in the testing phase, the bootstrapped Spearman correlation between the predicted feature and the actual f0 feature was calculated (median over 100 index-shuffles). To further validate the backward decoding results, we used a n-fold cross-validation approach, where in each fold a different section of the data is used for testing. The amount of folds n is equal to the amount of unique non-overlapping 2 minute sections of voiced data available for each story, i.e. 2 folds for Eline, 4 folds for DOL, and 3 folds for the other stories. The final backward correlation is the median correlation across these folds. The stronger the f0 response, the easier it is to train a high-quality model and the higher the backward correlations are. The backward correlations were compared to a story-specific significance level (based on correlations with spectrally-matched noise signals) to evaluate their statistical significance (two-sided test at α = 0.05). Story-specific differences between backward correlations were statistically evaluated in R (version 3.6.3., R Core Team (2018)), using linear mixed models (package lme4, version 1.1.21, Douglas et al. (2015)) with a random intercept per subject.

### 2.6. Forward modelling

In forward modelling, one attempts to reconstruct the data in each EEG channel based on a linear combination of the feature and time-lagged versions of the feature using the same ridge regression approach. In this case, time lags from −20 to 80 ms with 1/fs steps (fs = 1024 Hz) were taken into account. Forward modelling is less powerful than backward modelling since the information is not integrated across the scalp. Therefore forward models were calculated only for the subject-story combinations that yielded significant backward correlations. The weights of the optimal linear combination for each EEG channel form a temporal response function (TRF), which characterizes the linear transformation applied to the feature by the neural system. Single TRFs tend to be rather noisy and were therefore averaged over subjects and over a channel selection (see figure 4) (while ensuring consistent polarity).

One problem with forward modelling of the f0 feature is the large degree of autocorrelation, in both the response and the EEG, that occurs because the f0 stays relatively steady over multiple f0 periods. If, for example, the EEG response tracks the stimulus f0 with a neural delay of *t* then the cross-correlation of the actual EEG with the EEG predicted based on the f0 feature peaks at *t*, but also at 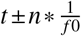 with lower peak amplitudes for larger n and alternating peak polarity. Accordingly, the TRFs will have large weights at each of these peaks. This hinders TRF interpretation as, instead of forming a single peak at *t*, the response activity is now spread out around *t* with both positive and negative peaks. Etard et al. (2019) proposed a method to aid in the interpretation of these TRFs and this method was also applied here (for code, see Kegler et al. (2018)): a Hilbert-transform was applied to the EEG data such that the forward model not only used time-shifted versions, but also phase-shifted versions of the EEG to predict the feature. This results in complex TRFs of which the absolute value was taken to obtain the ‘Hilbert’ TRF (hTRF). This discards the phase information and allows to focus on the amplitude variations in the hTRF. To evaluate at which latencies the hTRFs were significant, we determined a significance level (*alpha* = 0.05) based on forward modelling of mismatched data (more details in the appendix).

Apart from studying the temporal distribution of the responses using hTRFs, we also investigated the spatial distributions of the response at the peak latencies of the hTRFs. These spatial distributions, or topoplots, result from averaging the hTRF values at a specific lag over the selected subjects (with significant backward model) and plotting the result per channel on a head diagram with a color scale. A cluster-based permutation test from the mass-univariate ERP toolbox (Groppe et al., 2011) was applied to statistically evaluate the paired difference between two topoplots. A significance level of 0.05 was used and correction for multiple comparisons (64 channels) is implemented within the cluster test.

## 3. Results

### 3.1. Backward modelling

The correlations between the actual f0 and reconstructed f0 feature obtained with backward modelling are visualised for each story separately in figure 1. For the stories with a male voice, the correlations are significant for the majority of subjects (29 out of 35 for *Eline* and 33 out of 34 for *Milan*). In contrast, for the stories with a female voice, only a minority of the correlations exceed the significance level (7 significant data points for *AEDV*, 3 for *DKZ*, 3 for *DOL* and 7 for *DWZ*). None of the correlations are situated below the lower significance level, which is desired as negative correlations should not occur with reliable modelling.

To investigate whether story, speaker and voice gender of the speaker could predict the backward correlations, we applied a linear mixed model to the data (in R, package lme4, version 1.1.21, Douglas et al. (2015)) with story as a fixed effect and a random intercept per subject. Five contrasts were defined: 1) comparing stories with a female narrator and stories with a male narrator to evaluate the effect of voice gender (*AEDV, DKZ, DOL* & *DWZ* vs. *Eline* & *Milan*), 2) comparing stories with a different male narrator to evaluate speaker-specific differences for male voices (*Eline* vs. *Milan*), 3) comparing stories with a different female narrator to evaluate speaker-specific differences for female voices (*AEDV* vs. *DKZ, DOL* & *DWZ*) and 4) comparing stories with the same female narrator to evaluate story-specific differences (2 contrasts: *DKZ* vs. *DOL* and *DWZ* vs. *DOL*). Results indicate a significant difference between stories narrated by a female and a male voice (*β* = 0.059, df = 177, t = 22.04, p < 0.001). It is important to remark that this effect is caused by underlying acoustic differences in the voices (see further), and not the actual gender of the speaker. There is also a significant difference between *Eline* and *Milan*, stories with different male narrators (*β* = 0.026, df = 177, t = 5.78, p < 0.001). There was no significant difference between the stories with different female narrators (*β* = 0.0028, df = 177, t = 0.78, p = 0.44), nor was there a significant difference between the stories with the same female narrator (*DWZ* vs. *DOL*: *β* ∼ = 0, df = 177, t = 0.005, p = 0.99, *DKZ* vs. *DOL*: *β* = −0.0012, df = 177, t = −0.59, p = 0.56).

To further investigate the differences in response correlations across the stories, the correlations were related with two story parameters, i.e. the f0 and the rate of f0 change. The violin plots in figure 2 (left panel) show the f0 distribution for each story. As expected, the f0 is lower for stories with a male narrator than for stories with a female narrator. As can be seen in the right panel of figure 2, there is a negative linear relation between median f0 and backward correlation. A linear mixed model with a random intercept per subject confirms that this effect is significant (*β* = −0.00084, df = 177, t = −20.2, p < 0.001). Since the stories used in this study show a clear division in ‘low f0’ stories and ‘high f0’ stories, a group-wise comparison might be more adequate here: a linear mixed model with a random intercept per subject confirms that there is a significant difference between the ‘low f0’ stories and the ‘high f0’ stories (*β* = 0.059, df = 177, t = 20.2, p < 0.001).

In the left panel of figure 3, the distribution of the rate of f0 change of the stories is visualised. The median rate of f0 change is lower for male voices than for female voices. Among the stories with female narrator, *AEDV* had the lowest median rate of change. In the right panel of figure 3 the relation between the median rate of f0 change and the backward correlation (in percentage) is visualized. Higher median rate of f0 change is associated with lower backward correlations. A linear mixed model with a random intercept per subject confirms that this relation is significant (*β* = −0.00062, df = 178, t = −17.78, p < 0.001). When combining the effects of both median f0 and median rate of f0 change in one linear mixed model, both remain significant (median F0: *β* = −0.00064, df = 180, t = −6.27, p < 0.001, median rate of f0 change: *β* = −0.00017, df = 180, t = −6.27, p = 0.03).

**Figure 3:**
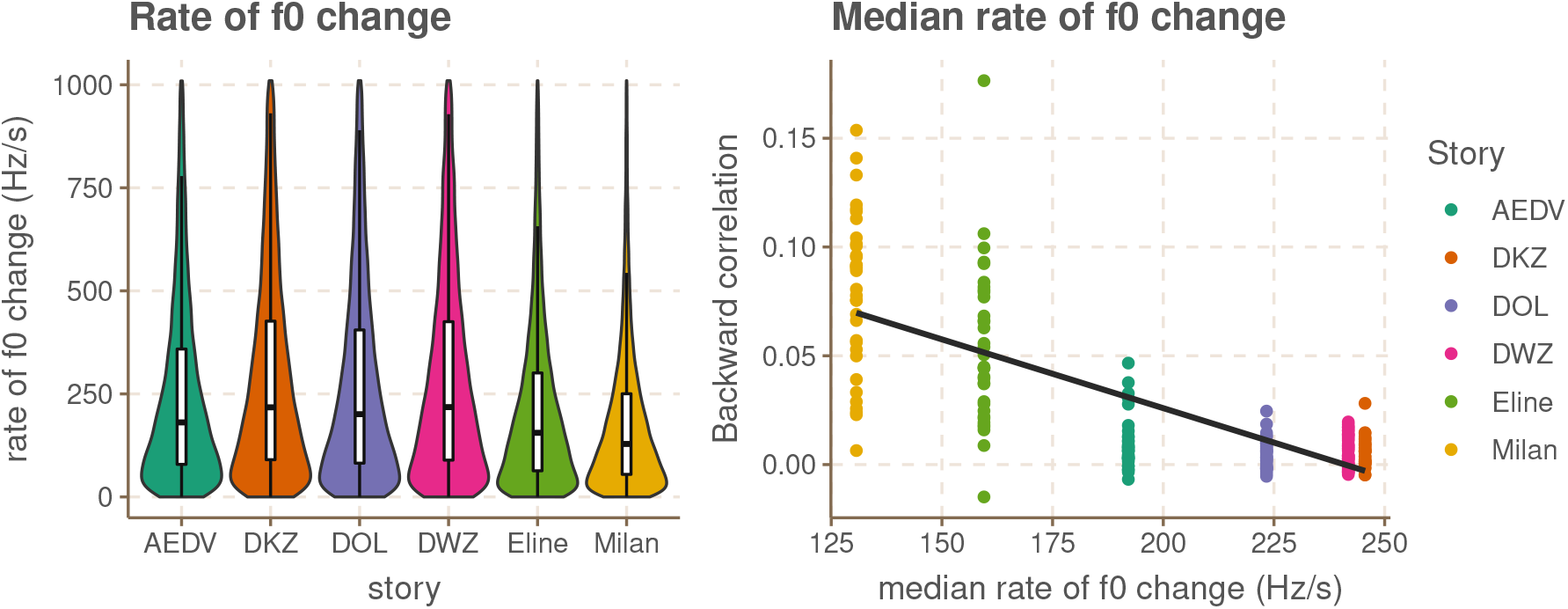
Rate of f0 change distribution per story and the relation between backward correlations and the median rate of f0 change of the story

**Figure 4:**
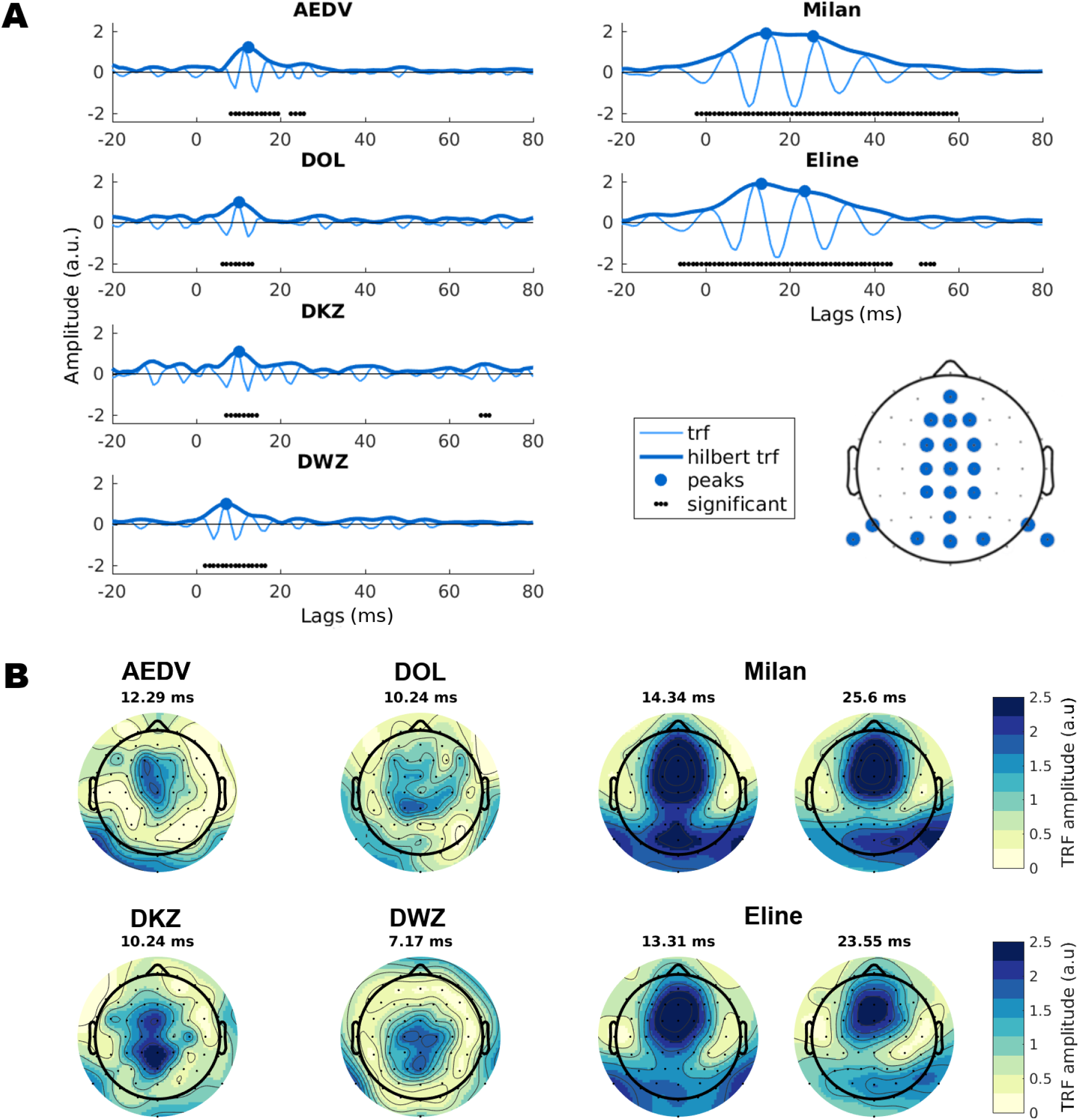
**A**. Average TRFs and hTRFS for female-narrated stories (left column) and male narrated stories (right column). TRFs and hTRFS were averaged over subjects with significant backward correlations and over the electrode selection indicated in the bottom right of the figure. Peaks in the hTRFs are indicated with a blue dot. Black dots at the bottom of the figure indicate lags (in milliseconds) where the hTRF is significantly different from noise. **B**. *Topoplots corresponding to hTRF peaks per story*. The hTRFs for female-narrated stories have a single peak and therefore a single topoplot. The hTRFs for male-narrated stories have two peaks and therefore two topoplots are shown.

### 3.2. Forward modelling

The final averaged TRFs and Hilbert TRFs (hTRFs) are visualised in panel A of Figure 4. As discussed before, the TRFs have a periodic nature due to the strong autocorrelation of the feature, and therefore also of the EEG response. This periodic nature makes it difficult to locate peak latencies that can be connected to neural sources. The hTRFs are helpful here because they do not display this periodic behaviour. However, we feel it is important to also display the regular TRF, as a reminder of the underlying process. The black dotted lines on figure 4 indicate lags where the hTRF significantly differed from noise (estimated through mismatched hTRFs). For the female narrated stories the significant latency range is located between 8.2 and 19.4 ms for *AEDV* (with a second smaller significant area between 22.5 and 25.6 ms), between 6.1 and 13.3 ms for *DOL*, between 7 and 14.3 ms for *DKZ* (with a second smaller significant area between 67.6 and 69.6 ms) and finally between 2 and 16.4 ms for *DWZ*. For the stories with male narrator the significant latency ranges are broader, in part because the autocorrelation effects cause broader spreading of activity for lower f0. Significant lags are located between −2 and 59 ms for *Milan* and between −6 and 44 ms for *Eline* (with a second significant area between 51.2 and 54.3). The stories with a female narrator gave rise to a hTRF with a single peak. The peak latency was equal to 12.29 ms, 10.24 ms, 10.24 ms and 7.17 ms, for the stories *AEDV, DOL, DKZ* and *DWZ* respectively. The hTRFs for stories with male narrator show peak activity along a broader range of latencies, which is harder to interpret. A first peak can be discerned at 13.31 ms for *Eline* and 14.34 ms for *Milan*, slightly later than the peaks observed for the stories with a female narrator. Furthermore, a second ‘peak’ can be discerned at later latencies, i.e. 25.6 ms for *Milan* and 23.55 ms for *Eline*. The female-narrated story AEDV seems to show a similar second peak around 25.6 ms which is small but significant.

The topoplots in panel B of figure 4 correspond to the peak latencies indicated on panel A. For the stories with a female narrator, the response activity at the peak latency is predominantly centrally located. For the stories with male narrator, the early response peak has strong frontocentral activity combined with bilateral occipito-parietal activity, which is stronger for *Milan* compared to *Eline*. At the later peak, the frontrocentral activity is slightly reduced and the parietal activity has become right dominant. A cluster-based permutation test on the paired difference between the topoplots corresponding to the earlier and later peaks revealed a significant difference for *Eline* (p = 0.017). The difference was not significant for *Milan* but the p-value is small (p = 0.09). The clustering that is formed for *Eline* indicates that the central activity (FC1, C1, C3, FCz, Cz, C2, CP2), the left mastoidal activity (TP7) and occipital (Oz) are decreased at the later latency. The non-significant clustering for *Milan* also indicates a decrease in central and occipital activity (CP1, P1, Pz, CPz). It should be noted that this statistical analysis suffers from the smearing of response activity across lags discussed earlier.

## 4. Discussion

In this study continuous EEG responses to the f0 were analysed for six stories with four different speakers using both forward and backward linear modelling. Average response strength was quantified through backward correlations. Story-specific differences between backward model correlations were related to the f0 of the voice and the rate of change of the f0 within each story. The TRFs and topoplots resulting from the forward models were studied to uncover spatial and temporal response patterns.

### 4.1. Backward correlations and the effect of stimulus parameters

Backward correlations between the reconstructed and the actual f0 feature were significantly larger for stories with a male narrator than for stories with a female narrator. Note that this effect is driven by typical intrinsic acoustic differences in female and male voices, not directly by the gender of the speaker. There was also a significant difference between the two stories with a male narrator, but not between the stories with the same or different female narrator (possibly partly because of a flooring effect). Further analysis indicated that these story/voice driven differences were largely linked to differences in f0 and rate of f0 change. Both of these stimulus factors were inversely related to the backward correlations, i.e. larger correlations for lower f0 and lower rate of f0 change. This observation is in line with the behaviour of classic EFRs which have a lower amplitude for higher stimulus frequencies and rapid frequency changes (Purcell et al., 2004; Billings et al., 2019; Van Canneyt et al., 2020). One major cause for this is that it is harder for neurons to phase-lock when the stimulus frequency is high and highly variable, resulting in weaker population responses.

We observed that for most of the subjects the correlations were not significant for the female narrated stories. An important reason for this is that female-narrated stories generally have higher f0 and also more variable f0 (higher rate-of-change) and that, as mentioned above, both of these factors are related to poorer backward correlations. Note that the absence of significant backward correlations, and by extension f0 responses, does not necessarily translate to poor pitch perception. Pitch encoding is thought to occur through rate-place code across large neural populations in the brainstem and higher up the auditory pathway (Micheyl et al., 2013), not purely temporal processing.

Analyzing EEG responses to the f0 in continuous speech with linear models is novel research that, to our knowledge, is only reported in one preceding study by Etard et al. (2019). The correlations we obtained for the male-narrated stories were in the range of what is reported in that study. However, Etard et al. (2019) also found significant correlations of around 0.05 with a female-narrated story for most of their subjects, which contrasts the findings of the present study. To understand this difference, we analysed the story with the female narrator used by Etard et al. (2019) and observed it had a median f0 of 166 Hz and a median rate of f0 change of 138 Hz/s. The median rate of change of the female-narrated stories in the present study were considerably higher than this (compare with Figure 3), which likely contributed to the lower correlations observed in this study. Moreover, the stimulus used by Etard et al. (2019) also had remarkably strong higher harmonics, a characteristic reminiscent of artificial speech. It has been shown that strong harmonic content leads to larger responses to f0 (Jeng et al., 2011; Laroche et al., 2013; Van Canneyt et al., 2020). To investigate whether these strong harmonics enhance the backward correlations for f0-tracking, we performed a pilot study (n = 3) comparing backward correlations for a naturally-spoken and speech-synthesized version of *DWZ*. Figure 5 depicts spectrograms of both versions and clearly shows the stronger harmonics in the speech-synthesized version. The rate of f0 change of the speech-synthesized stimulus is also somewhat lower. In the bottom right panel of Figure 5, it is shown that the backward correlations are considerably larger for the synthesized version of the story. Hence, we conclude that female-narrated stories can be successfully used for f0 responses to continuous speech but only if the rate of f0-change is low and the higher harmonics have large amplitude. Apart from using artificial speech, higher harmonics can be strengthened by high-pass filtering a speech stimulus to boost the higher frequencies. However, it is not guaranteed that findings from these specifically selected stimuli generalize to more typical female voices, since neural processing is clearly influenced by these unique stimulus properties.

**Figure 5:**
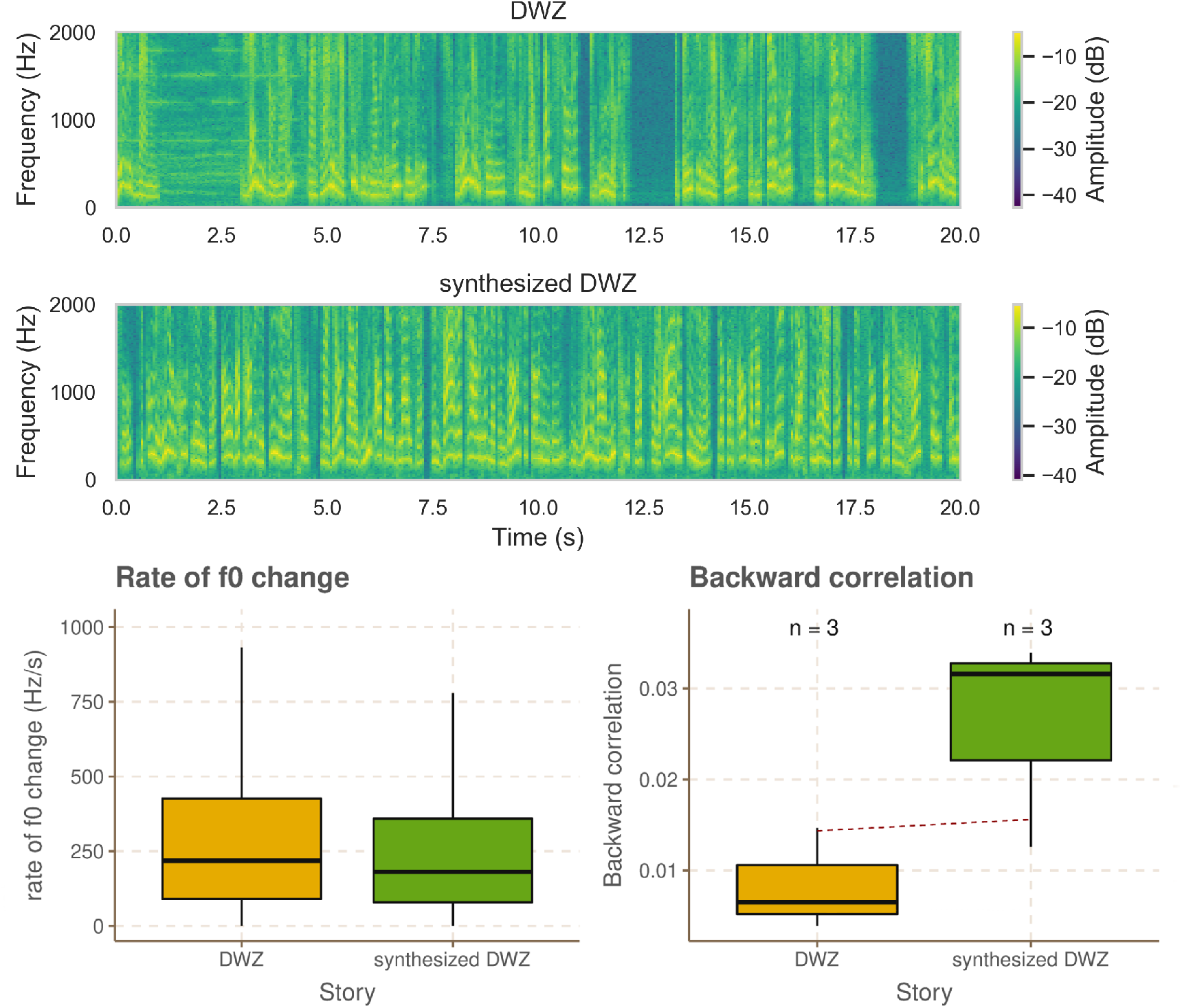
Spectrograms, rate of f0 change and backward correlations for the natural and speech-synthesized version of DWZ

### 4.2. Forward modelling and cortical contributions to the f0 response

Forward modelling allows analysis of the temporal and spatial response patterns through TRFs and topoplots. Importantly, forward modelling of f0 responses suffers from the large degree of autocorrelation in the f0 feature (and therefore also in the EEG). This leads to spreading of activation across lags in the TRFs, which makes them hard to interpret. The spreading is worse for lower f0 because the autocorrelation peaks are spread further apart. Moreover, the spreading is worse for lower rate of f0 change because the f0 is steady for more consecutive periods causing more and stronger autocorrelation peaks. Applying the Hilbert-technique of Etard et al. (2019) (hTRF) allows to take into account a π/2 phase-shift, which gets rid of the positive and negative deflections in the TRF, but does not remove the spread of activation across lags. Therefore the underlying TRF should be taken into account when interpreting the hTRF. Similar caution is necessary when interpreting topoplots as the spatial patterns are also smeared across lags.

TRFs reveal at which lags the dominant response activity occurs. The peak lag can then be related to a neural source, with longer lags corresponding to sources higher up the auditory pathway. When the hTRFs display a single peak, as is the case for most of the female-narrated stories in this study, the largest peak amplitude can easily be determined. For the responses to typical female voices, with high and highly variable f0, hTRF analysis indicated a peak-latency of 7-12 ms, which is similar to the latencies reported by Etard et al. (2019) and (Forte et al., 2017), and likely corresponds to a brainstem source (Langner and Schreiner, 1988). These results follow the well-researched notion that f0-evoked responses are predominantly generated by the inferior colliculus (Bidelman, 2015; Saiz-Alia and Reichenbach, 2020). It is important to note that these conclusions are drawn based on the limited amount of subjects who had significant backward correlations for the female narrated-stories.

For typical male-narrated stories, with lower and slowly varying f0, the hTRFs show broad activation which complicates the identification of peak lags as there could be multiple sources with different neural latency, whose activity is smeared together. In general, it is expected that the ‘smeared’ peaks will have lower amplitude and be symmetric around the main peaks. Therefore, patterns like what was observed for *Milan* and *Eline* suggest the presence of (at least) two sources with different latency. A first peak in the hTRF was identified at a latency of 13-14 ms, somewhat later than the peak latency observed for the female-narrated stories. However, (Bidelman, 2018) reported a highly similar peak latency for speech-evoked EFRs at 13.57 ms. Further, the male-narrated stories gave rise to an additional peak with a latency of 23.5-25.6 ms (which is perhaps also present for *AEDV*), suggesting contributions of an additional source when f0 is low and varies slowly. This additional source was not reported by (Etard et al., 2019) or (Bidelman, 2018), however, these results do closely the observations of (Kuwada et al., 2002) and Tichko and Skoe (2017) who both found EFR generators at 13 and 26 ms.

Topoplot analysis was done at the observed peak latencies to investigate which neural sources correspond to each of the responses peaks. Please note that the results from this analysis are tentative and need to be verified with detailed source analysis methods. For female-narrated stories, with variable and high f0, the topoplots revealed a central activation, consistent with the inferior colliculus, and a bilateral posterior temporal activation, possibly from contributions of the primary auditory nerve. This spatial pattern is in accordance with the results of Etard et al. (2019). For the male-narrated stories two topoplots at two different peak latencies, i.e. 13-14 ms and 23.5-25.6 ms, were compared. However, interpreting the difference was challenging because the response activity is ‘smeared’ over lags. At the first peak latency, i.e. 13-14 ms, the male-narrated stories revealed frontrocentral activity and bilateral posterior temporal activity. Based on the small latency and central activation pattern, dominant subcortical sources with primary auditory nerve contributions seem likely. However, Bidelman (2018) and Tichko and Skoe (2017) labelled the response activity at 13-14 ms as cortical, with Bidelman (2018) specifically identifying the primary auditory cortex. The second peak latency was considerably larger than latencies typically reported for a brainstem response and therefore points towards a source further-up the auditory pathway, in the thalamus or in the cortex. Visual comparison of the topoplots at the early and late latencies indicated that the second source is likely located posterior temporally, consistent with the primary auditory cortex, and this activation is right lateralised. This is in agreement with the results of Coffey et al. (2016, 2017) and Ross et al. (2020), who observed a right-lateralised cortical contribution from the primary auditory cortex to the EFR. Coffey et al. (2016) also show that the f0 response strength in the right primary auditory cortex (measured with MEG) was correlated with the musical experience as well as behavioural pitch perception performance. This is in line with the long-standing belief that the right primary auditory cortex specializes in pitch and tonal processing, e.g. Zatorre (1988) and Matsushita et al. (2015). The findings of this topoplot analysis add to the growing body of evidence that f0 responses can have cortical contributions (Coffey et al., 2016, 2017; Bidelman, 2018; Holmes et al., 2018; Ross et al., 2020), especially when f0 and rate of f0 change is low, as is the case for the typical male voice. In this light, the tradition to equate auditory responses to the f0 (either classic FFRs or continuous responses) with brainstem responses, needs to be revised. It is also reasonable to assume that cortical contributions play a part in the increased response strength for male voices.

The fact that cortical sources were less, or not at all, present for the female-narrated stories is likely explained by the fact that more central sources are less capable of phase-locking to higher frequencies and fast frequency changes, i.e. the higher up the auditory pathway the lower the maximum frequency that can be tracked. Bidelman (2018) estimated that cortical contributions produce up to 40 % of the EFR for frequencies around 100 Hz, but are completely absent for frequencies above 150 Hz. Therefore, stimulus frequency is a highly influential parameter that needs to be carefully considered when drawing conclusions in EFR research. For example, many studies have concluded that attention modulates subcortical processing because they observed attention effects on the EFR or on f0 responses (Hoormann et al., 2000; Galbraith et al., 2003; Lehmann and Schoönwiesner, 2014; Holmes et al., 2018; Etard et al., 2019). However, most of these studies have not taken into account the possibility of cortical contributions to the EFR or f0 response, as also argued by Hartmann and Weisz (2019). An important exception is the recent study by Holmes et al. (2018) who reported attentional modulation at lower (93-109 Hz) but not higher (217-233 Hz) frequencies, consistent with the presence of cortical contributions in the first but not in the second case.

## 5. Conclusion

Temporal processing of the f0 was studied for 6 different samples of continuous speech in young normal hearing volunteers. We used a novel method based on linear decoding and encoding models which allows to process continuous f0 responses. Response strength (measured through backward correlations) was larger for male-narrated stories compared to female-narrated stories, likely due to intrinsic differences in the voice characteristics. Response strength was predicted by the f0 of the voices as well as the rate of f0 change throughout the story. Voices with high f0 provided small responses, but a pilot test indicated that response strength can be boosted by selecting monotonous voices with strong higher harmonics, e.g. artificial speech. Spatio-temporal analysis revealed that the f0 responses were probably dominated by a central subcortical source in the case of voices with high and variable f0, but cortical sources (likely the right primary auditory cortex) can contribute for voices with low, steady f0. The above described findings from continuous f0 responses are in line with what is known from traditional event-related potentials. For future research, the use of continuous speech stimuli and the novel method are recommended as the evoked responses reflect speech processing throughout the auditory system in a quasi-natural listening scenario and the response strength as well as the spatio-temporal characteristics can be studied at each processing stage with the same data (Brodbeck and Simon, 2020).

## 6. Appendix

### detailed methods

#### 6.1. Regularization

Since estimating the backward and forward models is an ill-posed problem, complicated by linear dependency between EEG channels and their time shifted versions, regularization is necessary to obtain a single solution. This regularization was done using ridge regression (Tikhonov and Arsenin, 1977; Hastie et al., 2001; Machens et al., 2004) and the regularization parameter was estimated as the maximum value in the autocorrelation matrix of the EEG (+ time-shifted versions) for each subject. Sweeping over regularization parameters for a few subjects confirmed that this estimation of the regularization parameter was valid.

#### 6.2. Significance level of backward correlations

The obtained backward correlations were compared to a story-specific significance level to evaluate their statistical significance. The significance level was estimated by creating 1000 random Gaussian signals between 0 and 1 and filtering them to have the same spectrum as the actual f0 feature of each story. Then, the Spearman correlation between each of these spectrally-matched noise signals and the actual f0 feature is calculated. The 2.5th and 97.5th percentile of the resulting 1000 correlations respectively provide the lower (negative) and higher (positive) significance level. Significance levels were determined for each fold and the median over folds was taken to obtain the final significance levels for each story. Backward correlations that are smaller than the lower significance level or larger than the higher significance level were considered significantly different from correlations that occurs because of noisy data.

#### 6.3. Averaging over TRFs in forward modelling

Single TRFs tend to be rather noisy and were therefore averaged over subjects and over a channel selection. The channel selection was determined based on the topoplot analysis (see figure 4 panel B) and included a large frontocentral selection (AFz, Fz, F1, F2, FCz, FC1, FC2, CPz, Pz, C1, C2, CP1, CP2 and Cz) as well as some occipitoparietal electrodes (Oz, O1, O2, P9, P10, PO7 and PO8). Since TRFs for electrode channels on different sides of the head were averaged together, polarity changes were needed to ensure the TRF signals had the same polarity. This was done per story, by correlating all the TRFs with a reference TRF (manually selected from the first few TRFs) and flipping the polarity of the signal when the correlation coefficient was negative.

#### 6.4. The Hilbert TRF and its significance level

Etard et al. (2019) proposed a method to add phase-information to the encoding/decoding process and this method was applied here: the Hilbert transform was applied to the EEG and the forward model was used to predict both the real part of the Hilbert-transformed EEG and the imaginary part of the Hilbert-transformed EEG. The two sets of weights (for the real part and the imaginary part) were then combined to obtain complex TRFs. The same averaging and flipping operations were applied to these TRFs before the absolute value was taken to obtain the final ‘Hilbert’ TRF (hTRF). The hTRF allows to focus on the amplitude of the TRF by discarding the phase information.

To determine at which latencies the hTRFs were significant, we performed the same forward modelling process with a mismatch between the feature and the Hilbert-transformed EEG, i.e. the Hilbert-transformed EEG data for each subject was predicted based on 100 different mismatched features to obtain 100 mismatched hTRFs per subject (averaged over the same channel selection as described above) per story. This process was done for each story separately and only for subjects with significant backward models. The mismatched feature was constructed by randomly selecting continuous minute-long chunks from the feature and concatenating them to form a new feature. Since the real hTRFs were averaged over subjects, the same needs to be done with the mismatched hTRF. Therefore, for each subject, one hTRF was randomly selected out of the set of 100 available for that subject and these were averaged together to obtain the final averaged mismatched hTRF. This final averaging step was repeated 5000 times with random selection of hTRFs per subject in each iteration. The 5000 averaged mismatched hTRFs form a noise distribution, to which the actual hTRF can be compared. The critical value was chosen at a significance level of 0.05 Bonferroni-corrected by dividing by the amount of samples in the TRF, i.e. 104.

## 7. Acknowledgements

Authors would like to thank Bernd Accou and Wendy Verheijen for collecting the dataset used in this study. They were assisted in data collection by Amelie Algoet, Jolien Smeulders, Lore Kerkhofs, Sara Peeters, Merel Dillen, Ilham Gamgami en Amber Verhoeven. We also would like to acknowledge Marlies Gillis for collecting the pilot data presented in the discussion and thank Jonas Vanthornhout for his advice on the response analysis. This research was funded by TBM-project LUISTER (T002216N) from the Research Foundation Flanders (FWO) and also jointly by Cochlear Ltd. and Flanders Innovation & Entrepreneurship (formerly IWT), project 50432. Additionally, this project has received funding from the European Research Council under the European Unions Horizon 2020 research and innovation programme (grant agreement No. 637424, ERC starting grant to Tom Francart). The first author, Jana Van Canneyt, is supported by a PhD grant for Strategic Basic research by the Research Foundation Flanders (FWO), project number 1S83618N. Finally, the research is carried out with support from a Wellcome Trust Collaborative Award in Science RG91976 to Dr. Bob Carlyon and Jan Wouters, and with support from Flanders Innovation & Entrepreneurship through the VLAIO research grant HBC.2019.2373 with Cochlear. There are no conflicts of interest, financial, or otherwise.

## Abbreviations

AEDV: Anna en de vorst (story name)
DKZ: de kleine zeemeermin (story name)
DOL: de oude lantaarn (story name)
DWZ: de wilde zwanen (story name)
EEG: electroencephalogram
EFR: envelope following response
f0: fundamental frequency of the voice
hTRF: Hilbert-transformed temporal response function
TRF: temporal response function

## References

Accou, B., Monesi, M. J., Montoya, J., Van Hamme, H., and Francart, T. (2020). Modeling the relationship between acoustic stimulus and EEG with a dilated convolutional neural network. In 28th European Signal Processing Conference (EUSIPCO),, Amsterdam, Netherlands (in press).

Aiken, S. J. and Picton, T. W. (2008). Envelope and spectral frequency-following responses to vowel sounds. Hearing Research, 245(1-2):35–47.

American Clinical Neurophysiology Society (2006). Guideline 5: guidelines for standard electrode position nomenclature. Am. J. Electroneurodiagnostic Technol., 46:222–225.

Bidelman, G. M. (2015). Multichannel recordings of the human brainstem frequency-following response: Scalp topography, source generators, and distinctions from the transient ABR. Hearing Research, 323:68–80.

Bidelman, G. M. (2018). Subcortical sources dominate the neuroelectric auditory frequency-following response to speech. NeuroImage, 175(2018):56–69.

Billings, C. J., Bologna, W. J., Muralimanohar, R. K., Madsen, B. M., and Molis, M. R. (2019). Frequency following responses to tone glides: Effects of frequency extent, direction, and electrode montage. Hearing Research, 375:25–33.

Boersma, P. and Weenink, D. (2015). PRAAT: doing phonetics by computer.

Brodbeck, C., Presacco, A., and Simon, J. Z. (2018). Neural source dynamics of brain responses to continuous stimuli: Speech processing from acoustics to comprehension. NeuroImage, 172:162–174.

Brodbeck, C. and Simon, J. Z. (2020). Continuous speech processing. Current Opinion in Psychology, 18:25–31.

Broderick, M. P., Anderson, A. J., Di Liberto, G. M., Crosse, M. J., and Lalor, E. C. (2018). Electrophysiological Correlates of Semantic Dissimilarity Reflect the Comprehension of Natural, Narrative Speech. Current Biology, 28(5).

Coffey, E. B., Musacchia, G., and Zatorre, R. J. (2017). Cortical Correlates of the Auditory Frequency-Following and Onset Responses: EEG and fMRI Evidence. The Journal of Neuroscience, 37(4):830–838.

Coffey, E. B., Nicol, T., White-Schwoch, T., Chandrasekaran, B., Krizman, J., Skoe, E., Zatorre, R. J., and Kraus, N. (2019). Evolving perspectives on the sources of the frequency-following response. Nature Communications, 10(1):1–10.

Coffey, E. B. J., Herholz, S. C., Chepesiuk, A. M. P., Baillet, S., and Zatorre, R. J. (2016). Cortical contributions to the auditory frequency-following response revealed by MEG. Nature Communications, 7:11070.

Crosse, M. J., Di Liberto, G. M., Bednar, A., and Lalor, E. C. (2016). The multivariate temporal response function (mTRF) toolbox: A MATLAB toolbox for relating neural signals to continuous stimuli. Frontiers in Human Neuroscience, 10(NOV2016).

Decruy, L., Vanthornhout, J., and Francart, T. (2019). Evidence for enhanced neural tracking of the speech envelope underlying age-related speech-in-noise difficulties. Journal of Neurophysiology, 122(2):601–615.

Decruy, L., Vanthornhout, J., and Francart, T. (2020). Hearing impairment is associated with enhanced neural tracking of the speech envelope. Hearing Research, 393:107961.

Di Liberto, G. M., Crosse, M. J., and Lalor, E. C. (2018). Cortical measures of phoneme-level speech encoding correlate with the perceived clarity of natural speech. eNeuro, 5(2):1–13.

Di Liberto, G. M. and Lalor, E. C. (2017). Indexing cortical entrainment to natural speech at the phonemic level: Methodological considerations for applied research. Hearing Research, 348:70–77.

Di Liberto, G. M., O’Sullivan, J. A., and Lalor, E. C. (2015). Low-frequency cortical entrainment to speech reflects phoneme-level processing. Current Biology, 25(19):2457–2465.

Ding, N. and Simon, J. Z. (2012). Neural coding of continuous speech in auditory cortex during monaural and dichotic listening. Journal of Neurophysiology, 107(1):78–89.

Ding, N. and Simon, J. Z. (2014). Cortical entrainment to continuous speech: Functional roles and interpretations. Frontiers in Human Neuroscience, 8(MAY):1–7.

Douglas, B., Maechler, M., Bolker, B., and Walker, S. (2015). Fitting Linear Mixed-Effects Models Using lme4,. Journal of Statistical Software, 67(1):1–48.

Etard, O., Kegler, M., Braiman, C., Forte, A. E., and Reichenbach, T. (2019). Decoding of selective attention to continuous speech from the human auditory brainstem response. NeuroImage, 200(May):1–11.

Etard, O. and Reichenbach, T. (2019). Neural Speech Tracking in the Theta and in the Delta Frequency Band Differentially Encode Clarity and Comprehension of Speech in Noise. The Journal of neuroscience: the official journal of the Society for Neuroscience, 39(29):5750–5759.

Forte, A. E., Etard, O., and Reichenbach, T. (2017). The human auditory brainstem response to running speech reveals a subcortical mechanism for selective attention. eLife, 6:1–13.

Francart, T., van Wieringen, A., and Wouters, J. (2008). APEX 3: a multi-purpose test platform for auditory psychophysical experiments. Journal of Neuroscience Methods, 172(2):283–293.

Galbraith, G. C., Olfman, D. M., and Huffman, T. M. (2003). Selective attention affects human brain stem frequency-following response. NeuroReport, 14(5):735–738.

Groppe, D. M., Urbach, T. P., and Kutas, M. (2011). Mass univariate analysis of event-related brain potentials/fields I: A critical tutorial review. Psychophysiology, 48(12):1711–1725.

Hamilton, L. S. and Huth, A. G. (2020). The revolution will not be controlled: natural stimuli in speech neuroscience. Language, Cognition and Neuroscience, 35(5):573–582.

Hartmann, T. and Weisz, N. (2019). Auditory cortical generators of the Frequency Following Response are modulated by intermodal attention. NeuroImage, 203(June):116185.

Hastie, T., Tibshirani, R., and Friedman, J. (2001). The Elements of Statistical Learning. Springer, New York.

Haufe, S., Meinecke, F., Goörgen, K., Daähne, S., Haynes, J. D., Blankertz, B., and Bießmann, F. (2014). On the interpretation of weight vectors of linear models in multivariate neuroimaging. NeuroImage, 87:96–110.

Holmes, E., Purcell, D. W., Carlyon, R. P., Gockel, H. E., and Johnsrude, I. S. (2018). Attentional Modulation of Envelope-Following Responses at Lower (93-109 Hz) but Not Higher (217-233 Hz) Modulation Rates. JARO - Journal of the Association for Research in Otolaryngology, 19(1):83–97.

Hoormann, J., Falkenstein, M., and Hohnsbein, J. (2000). Early attention effects in human auditory-evoked potentials. Psychophysiology, 37(1):29– 42.

Jeng, F. C., Costilow, C. E., Stangherlin, D. P., and Lin, C. D. (2011). Relative power of harmonics in human frequency following responses associated with voice pitch in American and Chinese adults. Perceptual and Motor Skills, 113(1):67–86.

Kegler, M., Etard, O., Ae, F., and Reichenbach, T. (2018). Python code for the computation of complex TRFs (cTRFs). Github, (https://github.com/ReichenbachLab/cTRF.).

Krishnan, A., Xu, Y., Gandour, J. T., and Cariani, P. A. (2004). Human frequency-following response: Representation of pitch contours in Chinese tones. Hearing Research, 189(1-2):1–12.

Kuwada, S., Andersont, J. S., Batrat, R., Fitzpatrick, D. C., Teissier, N., and D ‘angelo, W. R. (2002). Sources of the Scalp-Recorded Amplitude-Modulation Following Response. J Am Acad Audiol, 13(2002):188–204.

Lalor, E. C. and Foxe, J. J. (2010). Neural responses to uninterrupted natural speech can be extracted with precise temporal resolution. European Journal of Neuroscience, 31(1):189–193.

Langner, G. and Schreiner, C. E. (1988). Periodicity coding in the inferior colliculus of the cat. I. Neuronal mechanisms. Journal of neurophysiology, 60(6):1799–1822.

Laroche, M., Dajani, H. R., Prévost, F., and Marcoux, A. M. (2013). Brainstem auditory responses to resolved and unresolved harmonics of a synthetic vowel in quiet and noise. Ear and Hearing, 34(1):63–74.

Lehmann, A. and Schoönwiesner, M. (2014). Selective attention modulates human auditory brainstem responses: Relative contributions of frequency and spatial cues. PLoS ONE, 9(1).

Lesenfants, D., Vanthornhout, J., Verschueren, E., Decruy, L., and Francart, T. (2019a). Predicting individual speech intelligibility from the cortical tracking of acoustic-and phonetic-level speech representations. Hearing Research, 380:1–9.

Lesenfants, D., Vanthornhout, J., Verschueren, E., and Francart, T. (2019b). Data-driven spatial filtering for improved measurement of cortical tracking of multiple representations of speech. Journal of neural engineering, 16(6).

Machens, C. K., Wehr, M. S., and Zador, A. M. (2004). Linearity of Cortical Receptive Fields Measured with Natural Sounds. Journal of Neuroscience, 24(5):1089–1100.

Matsushita, R., Andoh, J., and Zatorre, R. J. (2015). Polarity-specific transcranial direct current stimulation disrupts auditory pitch learning. Frontiers in Neuroscience, 9(MAY):174.

Mesgarani, N., David, S. V., Fritz, J. B., and Shamma, S. A. (2009). Influence of context and behavior on stimulus reconstruction from neural activity in primary auditory cortex. Journal of Neurophysiology, 102(6):3329–3339.

Micheyl, C., Schrater, P. R., and Oxenham, A. J. (2013). Auditory Frequency and Intensity Discrimination Explained Using a Cortical Population Rate Code. PLoS Computational Biology, 9(11).

Monesi, M. J., Accou, B., Montoya-Martinez, J., Francart, T., and Van hamme, H. (2020). An LSTM based architecture to relate speech stimulus to EEG. In ICASSP, IEEE International Conference on Acoustics, Speech and Signal Processing - Proceedings. IEEE.

Peng, F., Innes-Brown, H., McKay, C. M., Fallon, J. B., Zhou, Y., Wang, X., Hu, N., and Hou, W. (2018). Temporal Coding of Voice Pitch Contours in Mandarin Tones. Frontiers in Neural Circuits, 12(July):1–17.

Purcell, D. W., John, S. M., Schneider, B. A., and Picton, T. W. (2004). Human temporal auditory acuity as assessed by envelope following responses. The Journal of the Acoustical Society of America, 116(6):3581–3593.

R Core Team (2018). R: A Language and Environment for Statistical Computing. R Foundation for Statistical Computing, Vienna, Austria.

Ross, B., Tremblay, K. L., and Alain, C. (2020). Simultaneous EEG and MEG recordings reveal vocal pitch elicited cortical gamma oscillations in young and older adults. NeuroImage, 204.

Saiz-Alia, M. and Reichenbach, T. (2020). Computational modeling of the auditory brainstem response to continuous speech. Journal of Neural Engineering, in press:0–31.

Sohmer, H., Pratt, H., and Kinarti, R. (1977). Sources of frequency following responses (FFR) in man. Electroencephalography and Clinical Neurophysiology, 42(5):656–664.

Somers, B., Francart, T., and Bertrand, A. (2018). A generic EEG artifact removal algorithm based on the multi-channel Wiener filter. Journal of Neural Engineering, 15(3).

The MathWorks Inc. (2016). MATLAB: R2016b. Natick, Massachusetts.

Theunissen, F. E., Sen, K., and Doupe, A. J. (2000). Spectral-temporal receptive fields of nonlinear auditory neurons obtained using natural sounds. Journal of Neuroscience, 20(6):2315–2331.

Tichko, P. and Skoe, E. (2017). Frequency-dependent fine structure in the frequency-following response: The byproduct of multiple generators. Hearing Research, 348:1–15.

Tikhonov, A. N. and Arsenin, V. Y. (1977). Solutions of ill-posed problems. Scripta series in mathematics. V. H. Winston & Sons, Washington.

Van Canneyt, J., Hofmann, M., Wouters, J., and Francart, T. (2019). The effect of stimulus envelope shape on the auditory steady-state response. Hearing research, 380:22–34.

Van Canneyt, J., Wouters, J., and Francart, T. (2020). From modulated noise to natural speech: The effect of stimulus parameters on the envelope following response. Hearing Research, 393:107993.

Vanthornhout, J., Decruy, L., Wouters, J., Simon, J. Z., and Francart, T. (2018). Speech Intelligibility Predicted from Neural Entrainment of the Speech Envelope. JARO - Journal of the Association for Research in Otolaryngology, 19(2):181–191.

Verschueren, E., Somers, B., and Francart, T. (2019). Neural envelope tracking as a measure of speech understanding in cochlear implant users. Hearing Research, 373:23–31.

Zatorre, R. J. (1988). Pitch perception of complex tones and human temporal-lobe functiona. Journal of the Acoustical Society of America, 84(2):566–572.

